# High Content Phenotypic Profiling in Oesophageal Adenocarcinoma Identifies Selectively Active Pharmacological Classes of Drugs for Repurposing and Chemical Starting Points for Novel Drug Discovery

**DOI:** 10.1101/2020.01.20.912212

**Authors:** Rebecca E Hughes, Richard J R Elliott, Alison F Munro, Ashraff Makda, J Robert O’Neill, Ted Hupp, Neil O Carragher

## Abstract

Oesophageal adenocarcinoma (OAC) is a highly heterogeneous disease, dominated by large-scale genomic rearrangements and copy number alterations. Such characteristics have hampered conventional target-directed drug discovery and personalized medicine strategies contributing to poor outcomes for patients diagnosed with OAC. We describe the development and application of a phenotypic-led OAC drug discovery platform incorporating image-based, high-content cell profiling and associated image-informatics tools to classify drug mechanism-of-action (MoA). We applied a high-content Cell Painting assay to profile the phenotypic response of 19,555 compounds across a panel of six OAC cell lines representing the genetic heterogeneity of disease, a pre-neoplastic Barrett’s oesophagus line and a non-transformed squamous oesophageal line. We built an automated phenotypic screening and high-content image analysis pipeline to identify compounds that selectively modified the phenotype of OAC cell lines. We further trained a machine-learning model to predict the MoA of OAC selective compounds using phenotypic fingerprints from a library of reference compounds.

We identified a number of phenotypic clusters enriched with similar pharmacological classes e.g. Methotrexate and three other antimetabolites which are highly selective for OAC cell lines. We further identify a small number of hits from our diverse chemical library which show potent and selective activity for OAC cell lines and which do not cluster with the reference library of known MoA, indicating they may be selectively targeting novel oesophageal cancer biology. Our results demonstrate that our OAC phenotypic screening platform can identify existing pharmacological classes and novel compounds with selective activity for OAC cell phenotypes.

## Introduction

Combined, the two major histological subtypes of oesophageal adenocarcinoma (OAC) and oesophageal squamous cell carcinoma (OSCC), represent the sixth leading cause of cancer deaths worldwide with less than one in five patients surviving five years from diagnosis ^1^. A shift in epidemiology over the last 50 years has meant the incidence of OAC now vastly exceeds that of OSCC in western countries ^2^, accounting for more than 80 % of oesophageal cancers in the United States ^3^. Defining the optimal neoadjuvant treatment regime is an area of active investigation ^4^ as, current treatments all carry a significant risk of systemic toxicity, histological response rates remain poor ^5^ and only a limited subgroup of patients experience any survival benefit over surgery alone ^6,7^.

OAC is a highly heterogeneous disease, dominated by large scale genomic rearrangements and copy number alterations ^8^. This has made clinically meaningful subgroups and well validated therapeutic targets difficult to define. Clinical trials with new molecular targeted agents have predominantly been directed towards epidermal growth factor receptor (EGFR) and human epidermal growth factor receptor 2 (HER2) receptors ^9–12^ but thus far have proven unsuccessful. A potential explanation is the almost ubiquitous co-amplification of alternative receptor tyrosine kinases (RTKs) and downstream pathways leading to redundancy and drug resistance ^8,13,14^. An alternative to target based drug discovery, and increasing in popularity with technological advances, is phenotypic drug discovery, defined as the identification of novel compounds or other types of therapeutic agents with no prior knowledge of the drug target. Recent advances in phenotypic screening include automated high-content profiling ^15,16^. This approach involves quantifying a large number of morphological features from cell or small-model organism assays in an unbiased way to identify changes and phenotypes of interest. One benefit to this method is that a target does not need to be predefined but the mechanism-of-action (MoA) of hit compounds can be inferred by reference to known compound sets using multivariate statistics and machine learning approaches. Thus, this may prove a beneficial strategy for complex, heterogeneous diseases were target biology is poorly understood and where modern-target directed drug discovery strategies have made little impact on patient care, as exemplified by OAC.

Taking an unbiased, profiling approach to phenotypic screening, we chose to apply the Cell Painting assay to capture large amounts of information on cellular and subcellular morphology to quantify cellular state across a panel of genetically distinct OAC cell lines. Cell Painting is an assay developed to capture as many biologically relevant morphological features in a single assay so as not to constrain discovery to what we think we already know ^17,18^. Therefore, upon chemical perturbation we can detect changes in a subset of profiled features allowing a phenotypic fingerprint to be assigned to a particular perturbation or compound ^15,19–21^. These fingerprints can then be used to identify specific phenotypic changes of interest, identify compounds that cause strong alterations in cell morphology suggesting changes in cellular state or stress, or predict MoA by similarity comparison to reference libraries of well annotated compound mechanisms ^17,21^. However, this type of analysis is typically performed in a single ‘model’ cell line, chosen for its suitability to image analysis. As a proof of principle that high-content phenotypic profiling could be applied to a panel of morphologically distinct OAC and tissue matched control cell lines, we iteratively optimized cell culture conditions, cell plating densities and the Cell Painting assay staining protocol across our cell panel. Assay performance in terms of distinguishing distinct compound MoA for each cell type was evaluated by testing a small reference set of well annotated compounds representing eight distinct mechanistic classes and performing Principal Component analysis (PCA) and t-Distributed Stochastic Neighbor Embedding (t-SNE) to visualize clustering of distinct mechanistic classes. We further developed a machine-learning model capable of predicting MoA across the panel of heterogeneous OAC cell lines. Following assay validation we subsequently screened a library of 19,555 small molecules comprising target annotated probe compounds, approved drug libraries and two diverse chemical sets with unknown MoA. PCA clustering of compound fingerprints distinguished a number of phenotypic clusters composed of similar pharmacological classes active in the OAC cell lines. We also applied a Mahalanobis distance threshold and differential Z-score on our phenotypic data to identify compounds from our screen which were selectively active in OAC versus tissue matched control cells. For prioritized hits we have selected a subset and validated OAC selectivity with follow up dose-response testing and performed transcriptomic pathway analysis pre- and post-treatment on sensitive and insensitive cell lines to further elucidate MoA. We further applied PCA and machine learning analysis to phenotypic fingerprints from our diverse chemical set to identify compounds which exhibit selective activity upon OAC cell phenotypes by a mechanism distinct from our reference set indicating they may exhibit novel MoA.

Herein we describe the development and validation of a high-content phenotypic profiling assay and associated image-informatics and machine learning toolbox to classify MoA of phenotypic screening hits across a panel of OAC and tissue-matched control cell lines. This approach has enabled the identification of chemical and target classes, including HDAC inhibitors, which consistently cause the same cellular response across the panel of OAC lines, demonstrating efficacy against the heterogeneity of the disease. In addition, we identify pharmacological classes such as the antimetabolites and new chemical entities with high selectivity for some OAC cell lines relative to tissue matched controls. We propose that applying high-content multiparametric phenotypic profiling to a panel of genetically annotated OAC cell lines may stimulate new drug discovery and drug development programs for OAC by the identification of drug repurposing opportunities and novel chemical starting points with selective activity for specific OAC genotypes.

## Materials and Methods

### Cell Culture

EPC2-hTERT cells were a kind donation from Anil Rustgis’ Lab, University of Pennsylvania ^22^.

### Cell Line Authentication

Cell line identification (not carried out for the EPC2-hTERT line, as there is no reference sequence) was confirmed by short tandem repeat (STR) genotyping (Cell Line Authentication, Public Health England).

The cell-lines were confirmed to be mycoplasma negative using the Venor™GeM Mycoplasma Detection PCR kit (MP0025; Sigma).

### Cell Subculture

Oesophageal adenocarcinoma (OAC) lines were grown in RPMI (#31870025, Life Technologies) supplemented with FBS (10 %) and L-glutamine (2 mM) and incubated under standard tissue culture conditions (37 °C and 5 % CO_2_). The Barrett’s oesophagus line; CP-A, and the oesophageal epithelial line; EPC2-hTERT, were grown in KSFM (#17005075, Gibco) supplemented with human recombinant EGF (5g/L) and BPE (50 mg/L). Soybean trypsin inhibitor (250 mg/L, 5 mL) was used to neutralise trypsin.

### High-Content OAC Cell Painting Assay

Cells were seeded (50 µL per well) into 384-well, CELLSTAR® Cell Culture Microplates (#781091, Greiner), and incubated under standard tissue culture conditions for 24 hours before the addition of compounds. CP-A cells were seeded at 800 cells per well, SK-GT-4 cells were seeded at 1000 cells per well and the remaining cell lines were all seeded at 1500 cells per well.

Compound source plates were made at 1,000-fold assay concentration and added to the cells with an overall dilution in media of 1:1000 from source to assay plate. Library concentrations (**Supplementary Table S1**).

After 48 hours incubation in the presence of the compounds, cells were fixed by the addition of an equal volume of formaldehyde (8 %, 50 µL) (#28908, Thermo Scientific) to the existing media, incubated at room temperature (20 minutes) and washed twice in PBS. Cells were then permeabilised in Triton-X100 (0.1%, 50 uL) and incubated at room temperature (20 minutes) followed by two more washes with PBS.

The staining solution (**Table 1**) was prepared in bovine serum albumin solution (1 %). Staining solution was added to each well (25 µL) and incubated in the dark at room temperature (30 minutes), followed by three washes with PBS and no final aspiration. Plates were foil sealed.

**Table 1:**
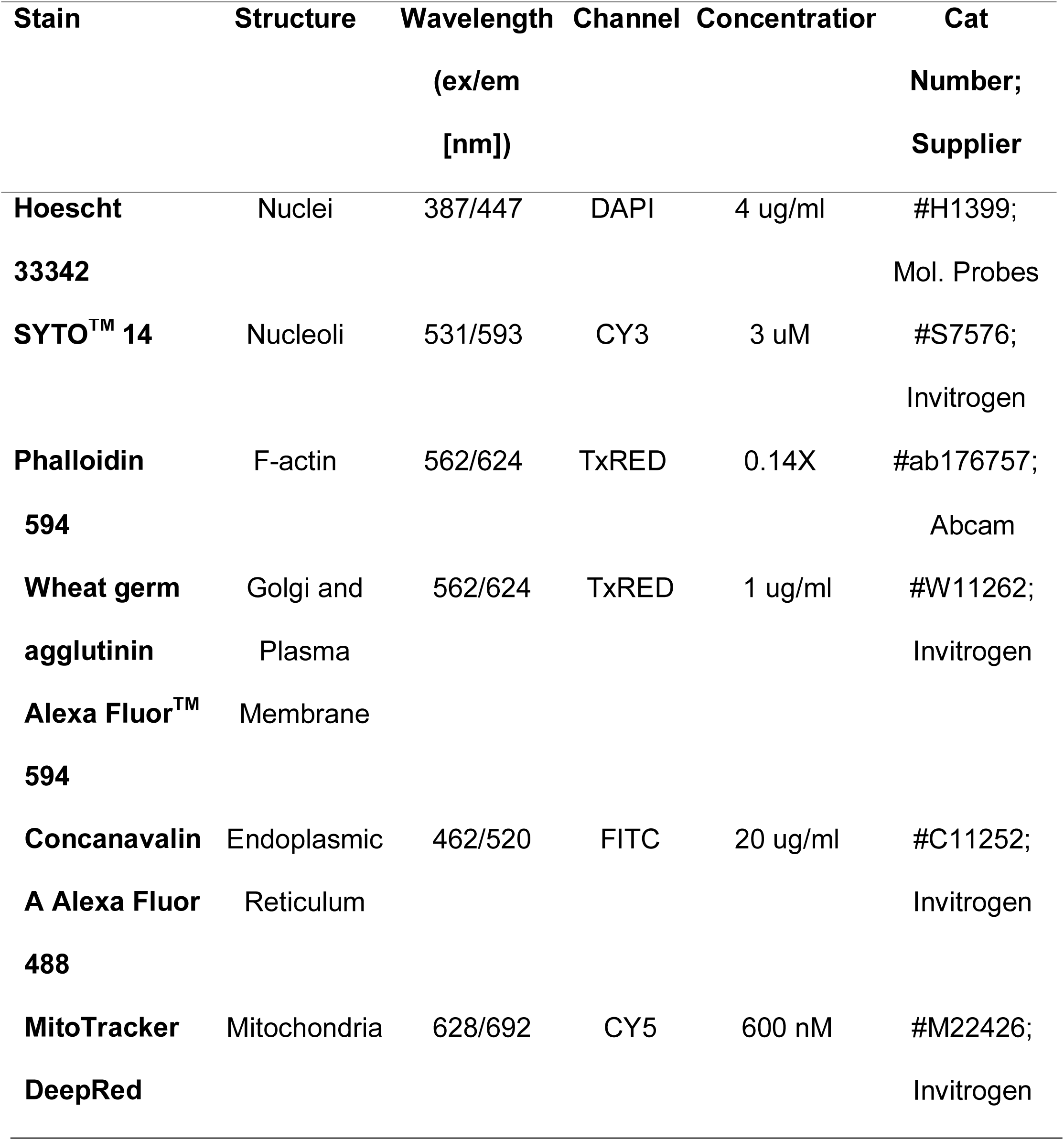
Cell painting reagents, concentrations, excitation/emission wavelengths of the filters used for imaging, and suppliers. ex: excitation, em: emission

### Image Acquisition

Plates were imaged on an ImageXpress micro XLS (Molecular Devices, USA) equipped with a robotic plate loader (Scara4, PAA, UK). Four fields of view were captured per well using a 20x objective and five filters (**Table 1**).

### Image Analysis

#### CellProfiler 2D image analysis

CellProfiler v3.0.0 ^23^ image analysis software was used to segment the cells and extract 733 features per cell per image. First the pipeline identified the nuclei from the DAPI channel and used these as seeds to aid a segmentation algorithm to identify the cell boundaries from the TxRed channel, and finally these two masks were subtracted to give the cytoplasm. These three masks marking the cellular boundaries were then used to measure morphological features including size, shape, texture, and intensity per object across the five image channels.

#### Image Preprocessing

The cell level data was aggregated to image level by taking the median for each measured feature per image. Low quality images and image artefacts were then identified and removed using image quality metrics extracted by CellProfiler. Images with less than 20 cells in them were also removed from final analysis. For the remaining images, features were normalised on a plate-by-plate basis by dividing each feature by the median DMSO response for that feature. Features with NA values were removed, as were features with zero or near zero variance, using the findCorrelation and nearZero functions in the R package Caret. All remaining features were scaled and centred globally by dividing by the standard deviation of each feature and subtracting the feature mean respectively. The pair-wise correlations were calculated for all remaining features, and highly correlated features (>0.95) were removed. Finally, the image level data was aggregated to the well (compound) level and this was used in the analysis.

#### Random Forest Classifier

The random forest classifier was implemented using R’s Random Forest package with the following specified parameters; ntree = 500, data was stratified by class and the sample size was set to the smallest class size for balance. The images from three concentrations for each compound were pooled and treated as a single class. Two different analyses were run, firstly MoA prediction was implemented for each cell line individually, and then using leave one out (LOO) cross-validation, leaving one OAC cell line out of the training set at a time and running that line as a test set.

Principal component analysis and T-SNE were implemented using built-in R functions, prcomp and RTSNE respectively to visualise the clustering of the compounds for each cell line.

#### Hierarchical Clustering

Z-scores and Mahalanobis scores were centred and scaled for each compound across the panel of cell lines. Spearman correlation was then used to generate a distance matrix and hierarchical clustering was determined using complete linkage.

#### NanoString Transcriptomic Analysis

Cells were seeded in 6-well plates and incubated for 24 hrs. Media was then removed and replaced with DMSO (0.1%) or Methotrexate (5 µM) in DMSO and incubated for 6 hours. Cells were scraped and lysed using QIAshredders (#79654, QIAGEN), RNA was extracted by means of the Qiagen RNeasy Mini kit (#74104, QIAGEN) (with β-mercaptoethanol) according to manufacturer instructions, and included a DNase digestion step (#79254, QIAGEN).

Of the purified RNA, 100□ng were used as input for amplification-free RNA quantification by the NanoString nCounter Analysis System with the Human PanCancer Pathways and Metabolic Pathways panels. Raw counts were normalised to the internal positive controls and housekeeping genes, using the nSolver 4.0 software.

## Results

### Assay development

Since OAC is such a heterogeneous disease, we chose to develop a high-content phenotypic screening assay composed of a panel of OAC and tissue matched non-transformed cell lines that captured this heterogeneity and thus provides a discovery platform for identification of novel targets and drug MoA which selectively target OAC. We assessed the amenability of 12 cell lines to high-content profiling, 10 OAC lines (JH-EsoAD1, FLO-1, MFD-1, OE33, OACM5.1, OAC-P4C, SK-GT-4, ESO51, ESO26 and OE19), and two tissue matched non-transformed lines; a Barrett’s oesophagus line CP-A, and a normal oesophageal squamous line immortalized by expression of telomerase EPC2-hTERT. We assessed each cell line against a criteria list that indicated high performance for high-content screening including, cell adhesion quality, cellular morphology, proliferation in 384-well plates, image segmentation, and MoA prediction accuracy. These criteria ensure image quality/information content, high-throughput screening compatibility and image segmentation accuracy for downstream analysis pipelines. Based on suitable cell adhesion and morphological properties we took forward the following eight cell lines for high-content assay development including image segmentation and machine learning analysis (CP-A, EPC2-hTERT, FLO-1, JH-EsoAD1, MFD-1, OAC-P4C, OE33, and SK-GT-4) (**Figure 1**).

**Figure 1.**
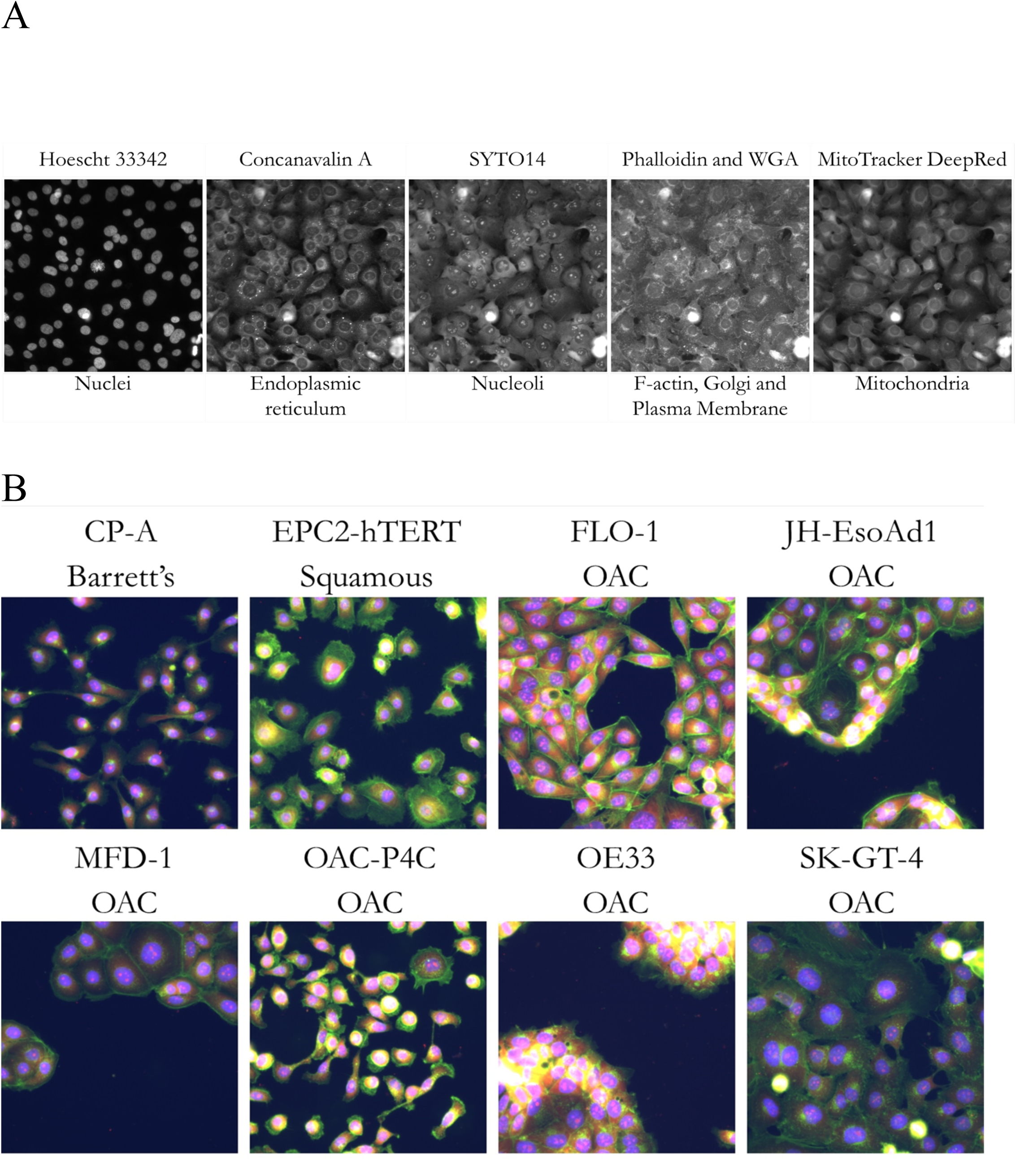
Cell Painting Assay and the OAC Cell Panel. **A)** The five channels imaged in the Cell Painting assay for representative cell line FLO-1, with dyes and cellular structures labelled. **B)** Colour combined representative control (DMSO) images of the six cell lines in the cell panel: DAPI (blue), TxRED (green), CY3 (red). See Table 1 for additional details about the stains and channels imaged.

The published Cell Painting protocol ^17,18^ was adapted for our cell lines specifically, as follows; the MitoTracker DeepRed was originally added before the cells were fixed, however, morphological changes have been seen in certain cell lines upon the addition of MitoTracker. Therefore we opted to fix the cells first and add all of the Cell Painting reagents together post-fixation to prevent artefactual morphological changes due to cell staining, and to reduce complexity for robotic handling in a high-throughput setting. This also necessitated that we re-optimise the dye concentrations across our cell panel. Here we increased the MitoTracker DeepRed concentration and reduced the concentration of Hoechst, Concanavalin A, and Wheat Germ Agglutinin and switched to a different phalloidin supply (**Table 1**).

### Machine learning

Standard assay quality control metrics such as Z’Factor are unsuitable for multiparametric assays, particularly cell based phenotypic profiling assays where a desired phenotype is unknown and/or there is a lack of positive controls ^24–26^. In order to assess assay quality from a compound MoA profiling perspective we used MoA prediction accuracy on a small well-annotated reference library of compounds with well-defined, known MoA (**Supplementary Table S2**). For this, we trained a random forest classifier using the CellProfiler extracted phenotypic information from the images of cells treated with the reference set of compounds.

Accuracy in the ability to predict MoA was used to assess whether, the OAC and tissue matched control cell lines were amenable to the phenotypic profiling assay, further validate image segmentation was accurate, and ensure that the phenotypic information extracted was relevant and broad enough to allow accurate prediction of MoA. In order to robustly evaluate compound selectivity and MoA across our heterogenous panel of genetically distinct OAC cells it was particularly important to assess the performance of each individual cell line and ensure that one cell line did not perform significantly better or worse than the others. A characteristic of OAC cell lines (OE33, MFD-1 and SK-GT-4 in particular), is the migration and formation of cell clumps, which are challenging to segment accurately by automated image analysis. Here we wanted to confirm that they were equal to the rest of the panel and suitable for the assay pipeline. OAC-P4C is a particularly morphologically heterogeneous line so it was also important to ensure that image level data can be used for phenotypic compound profiling in these types of cell lines.

In order to visualise the phenotypic information extracted, we performed two data reduction methods; PCA and T-SNE, on the well level data for the small reference library of compounds and plotted the first two components, coloured by mechanistic class. These results demonstrate that distinct compound classes (e.g. HDAC inhibitors) generally cluster together, however some classes such as statins do not produce strong phenotypes and clustering shows they are close to the DMSO controls, data for FLO-1 and MFD-1 cells are provided as exemplars. (**Figure 2**).

**Figure 2.**
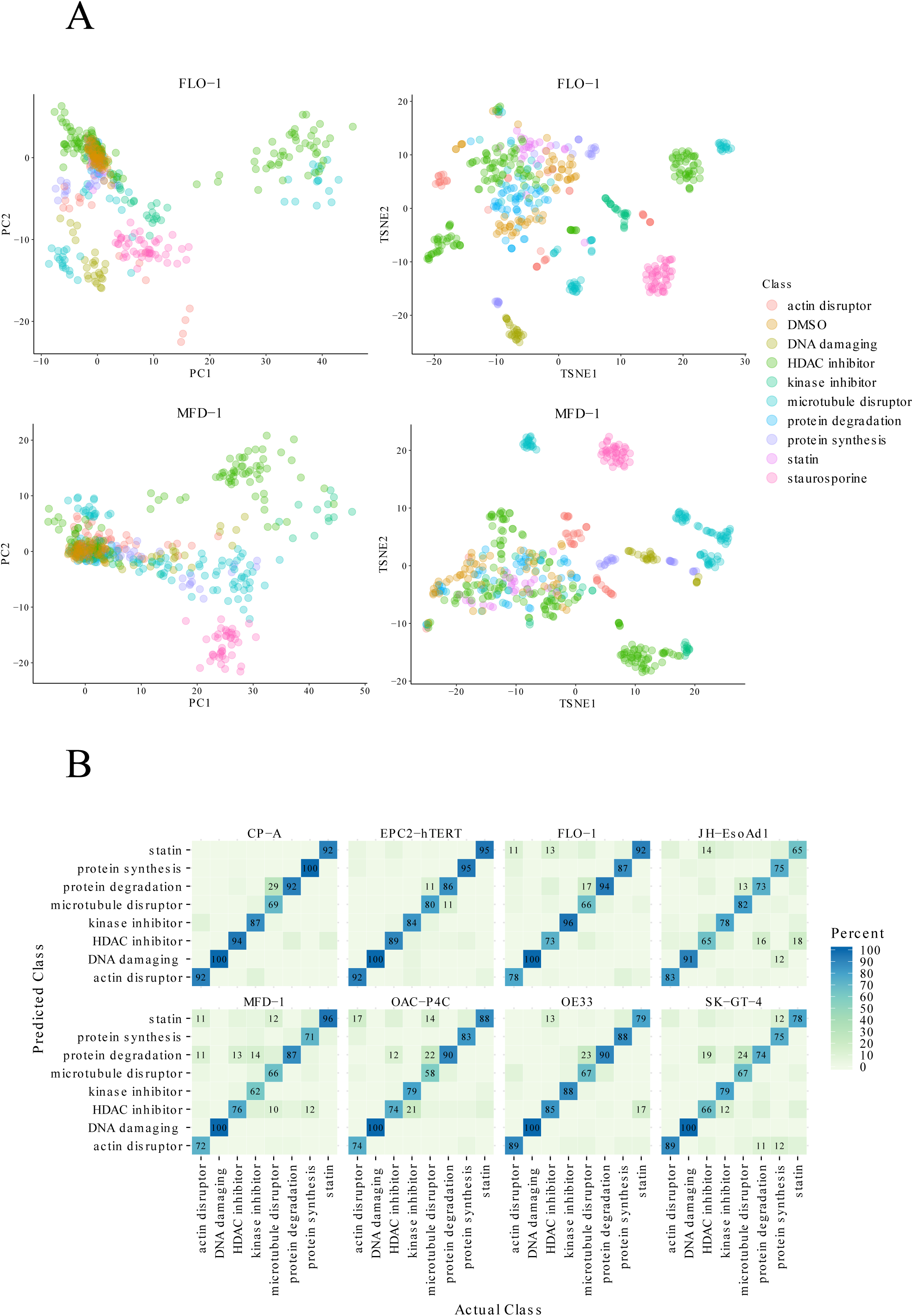
Reference Library Clustering and Machine Learning. **A)** The first two components of principal component analysis (PCA) and t-distributed stochastic neighbour embedding (T-SNE) for the reference library compound treatments for the OAC line FLO-1 and the Barrett’s line CP-A. Points are coloured by mechanistic class and multiple compounds concentrations are plotted. **B)** Random Forest Classifier: Confusion Matrices of prediction accuracies per cell line in the cell panel for the reference library of compounds. Diagonal values show class sensitivities.

We next optimised a random forest classifier to test MoA prediction on our reference library of well annotated compounds. The extracted features from three concentrations of each compound were pooled and used to train the classifier. We chose 0.1, 1 and 10 µM, as using a broad range of concentrations means that each compound does not need optimising individually across each cell line.

When trained and tested on each individual cell line the average out-of-bag error was 20.38 % across the entire panel of cell lines. The variation between cell lines (12-27 %) was also low, with no cell line dropping below 70 % accuracy, demonstrating the assay was well optimised across the panel. The weakest cell line was the OAC-P4C, which can likely be attributed to its heterogeneous morphology.

In order to confirm that the classifier was not overfitting we used leave-one-out cross-validation. We implemented leave-one-cell-line-out and trained it on five of the OAC lines, testing on the remaining line. Here, as expected it performed less well overall (**Supplementary Fig. S1**). However, there is still a very strong diagonal trend in the confusion matrices indicating the ability of this classifier to be transferred to new cell lines despite having no prior training on them and thus potential for application of the classifier across a broader panel of cell lines without the need to train each cell line individually.

Overall the accuracy of the machine learning demonstrates that the phenotypic profiling assay is of high quality across all eight cell lines, including morphologically heterogeneous cells, and feature extraction produces meaningful data for phenotypic analysis. The phenotypic profiling assay can therefore be applied to provide an initial evaluation of MoA of hit compounds influencing OAC cell proliferation, survival, and morphology. As such our multiparametric high-content phenotypic profiling assay may prove useful in prioritization of compound hits which represent novel MoA and de-prioritization of compound which represent undesirable MoA for subsequent medicinal chemistry and target deconvolution investments. We therefore prioritised the full panel of eight lines that passed our quality control criteria; six OAC lines with diverse genetic backgrounds, a Barrett’s oesophagus line, and a non-transformed squamous oesophageal line, for a phenotypic screen of 19,555 small molecules.

### Small molecule screen

A total of 19,555 small molecules including approved drugs were profiled against our panel of eight cell lines using the ImageXpress microXL high-content imaging platform. Cells were treated with the commercially available Prestwick Chemical Library of 1280 mostly off-patent drugs, the LOPAC library of pharmacologically active compounds (1280 compounds), a proprietary diverse chemical library provided by CRUK therapeutics discovery laboratories (13408 compounds), the BioAscent library of 3,200 compounds and bespoke libraries of 387 target-annotated compounds and chemical probes. The primary phenotypic screen across all eight cell lines encompassed, 512×384 well plates, 3.9 million images and 36 TB of data in total. Image analysis was performed using CellProfiler across a computer cluster.

Using a panel of cell lines better represents a heterogeneous disease, and allowed us to identify compounds which demonstrated selective activity across multiple OAC lines and not in the tissue matched control. We ran two parallel analyses for primary hit selection against the OAC lines; one based on broad, morphological, phenotypic changes and the other on cell growth and survival using nuclei count. At cytotoxic concentrations there are few attached cells and these are often rounded up leading to a lack of information in the images. Therefore, images with 20 cells or less were removed from the morphological analysis.

The following results focus on a subset of 3000 annotated compounds (excluding the CRUK Therapeutics labs and BioAscent lead like molecules).

In order to identify compounds inducing strong phenotypic changes we used PCA on the feature data to reduce the dimensions and then calculated the Mahalanobis distance to the DMSO controls for the first 15 principal components, which explain approximately 90% of the variation in the data across each cell line. The Mahalanobis distance therefore provides an unbiased metric of compound activity upon each cell line in the screen.

Phenotypic analysis identified 62 compounds that selectively target two or more of the OAC lines over the non-transformed oesophageal cells. Clustering the cell panel’s responses to these molecules showed a number of phenotypic clusters enriched with similar pharmacological classes, including HDAC inhibitors, microtubule disruptors, and antimetabolites, suggesting that hits have clustered mechanistically (**Figure 3A, Supplementary Fig. S2**).

**Figure 3.**
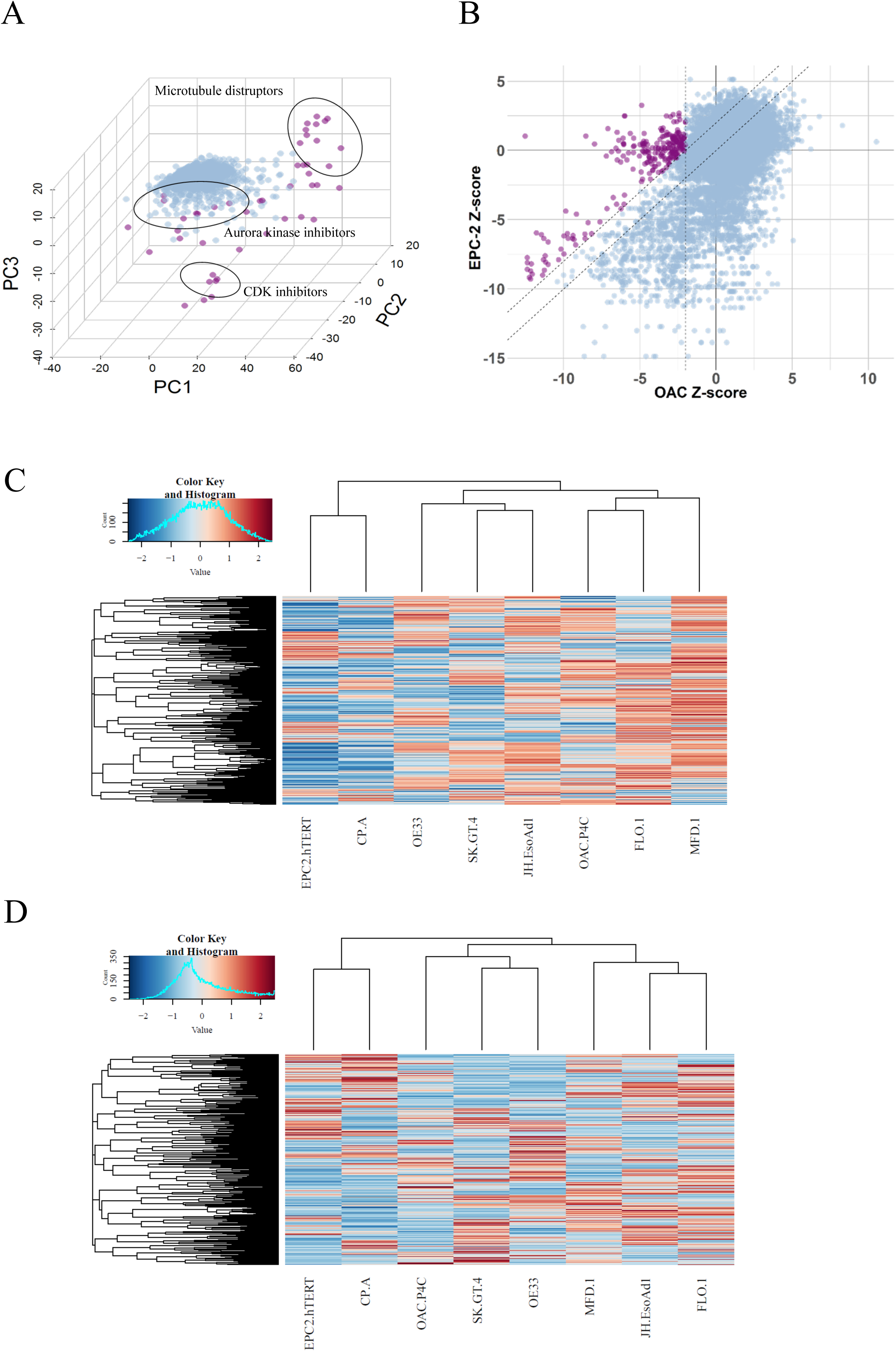
Hit analysis. **A)** The first three components of principal component analysis (PCA) for exemplar data from OAC cell line; JH-EsoAD1. Hits (purple) are defined as having a Mahalanobis distance of greater than 1500 from the DMSO controls. **B)** Z-score plot for all OAC lines overlaid vs EPC2-hTERT oesophageal squamous control line. Hits (purple) are defined as having a z-score of – 3 or more in the OAC lines and showing selectivity of at least 2 z-scores compared to the EPC2-hTERT line. **C)** Z-score hierarchical clustering of the cell panels response to compounds. **D)** Mahalanobis distance clustering of phenotypic response to compound treatments across cell lines.

Based on cell growth and survival (i.e nuclei count), we identified 27 compounds that were selectively active in two or more of our OAC lines. Here, hits were defined as having a z-score of −3 or more in the OAC lines and a difference of at least 2 in one or both of the control cell lines e.g. for a hit with a z-score of −3 in an OAC line, the z-score in the EPC-2 would have to be greater than or equal to 0. This comparison was made between each OAC line and the control lines to define hits and then selected if they were selectively active in at least two OAC lines across the panel (**Figure 3B**).

Compounds from the growth and survival analysis cluster into several therapeutic classes suggesting mechanistic pathways that may be selective for OAC cell growth and survival. Classes include antimetabolites and HDAC inhibitors. These classes were also identified in the morphometric phenotypic analysis (**Supplementary Fig. S2**).

We performed hierarchical clustering of cell line responses to the compounds, as determined by the Mahalanobis metric (morphometric phenotypic analysis) and the z-scores (nuclei count) (**Figure 3C and D**), enabling pharmacological discrimination of cell lines. These results show that the control cell lines; EPC2-hTERT and CP-A, can be discriminated from the OAC panel based on global drug screening data, providing confidence that our high-content cell painting assay can identify compounds with selectivity for OAC over the tissue matched control lines.

### Antimetabolites are selectively lethal to OAC cells

From the subset of 3000 annotated compounds we identified the drug Methotrexate and three other structurally related antimetabolites; Pemetrexed, Raltitrexed and Aminopterin, as highly selective for OAC cell lines relative to tissue matched control CP-A and EPC2-hTERT cells in both the nuclei count and morphological phenotypic analyses. We therefore validated this class of compound for dose dependent activity. Aminopterin was removed from further analysis due to its toxicity profile in the clinic ^27^, however, it showed potent activity in an initial dose response in the OAC lines, validating it as a hit our screen (results not shown).

Nuclei count dose responses for Methotrexate, Pemetrexed and Raltitrexed demonstrated strong selectivity against the OAC lines, with IC50s ranging from 1-65 nM, and showed minimal cytotoxic or phenotypic activity in either the CP-A or the EPC2-hTERT line even at 10 µM (**Figure 4A, Supplementary Table S3**), validating our hit selection criteria.

**Figure 4.**
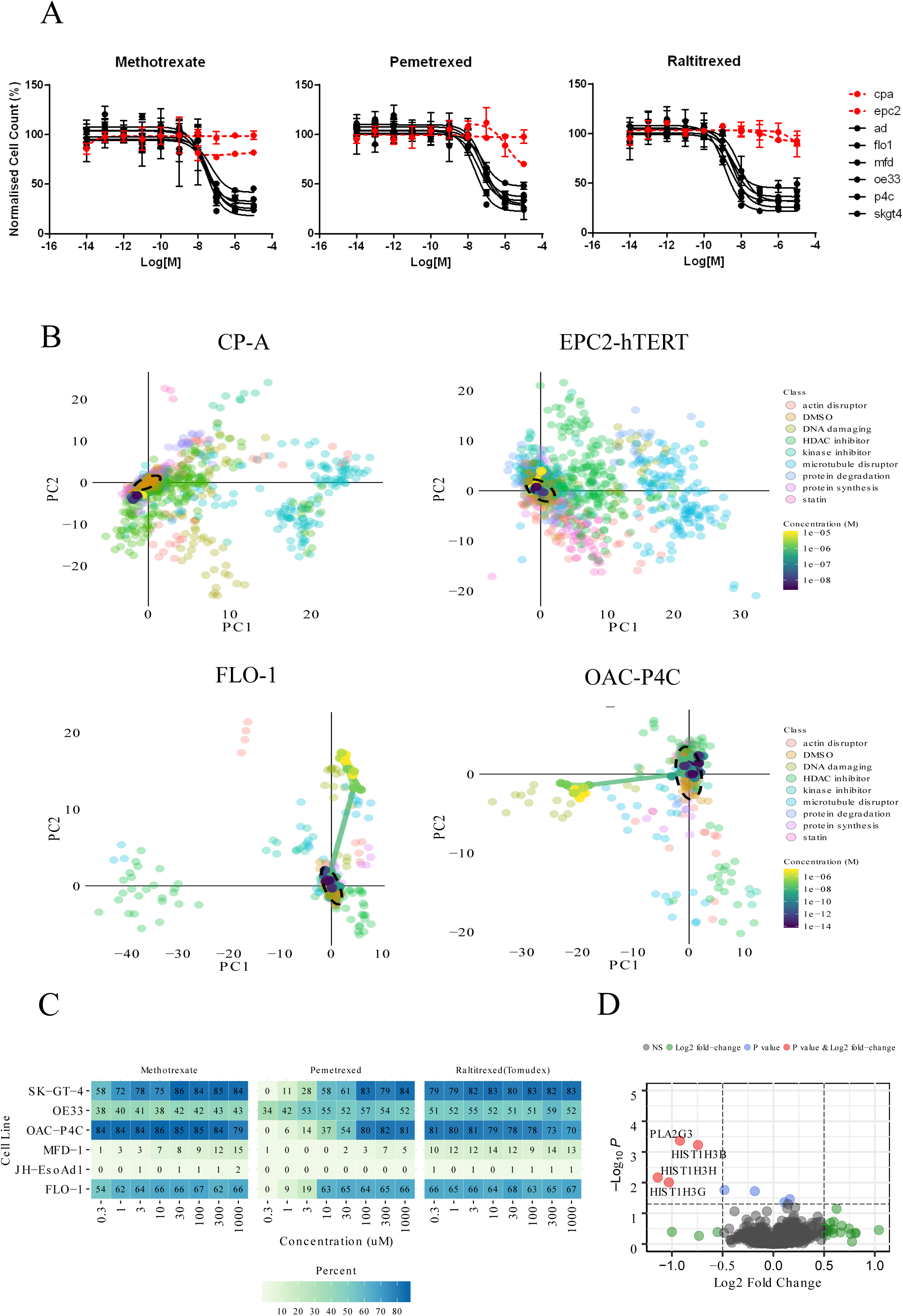
Antimetabolite evaluation. **A)** Dose responses for Methotrexate, Pemetrexed, and Raltitrexed across panel of cell lines. **B)** Principal component analysis of dose responses overlaid on reference library for Methotrexate in two resistant lines (CP-A and EPC2-hTERT) and two sensitive lines (FLO-1 and OAC-P4C). **C)** Probabilities expressed as percentages for DNA damaging class for each cell line and each of Methotrexate, Pemetrexed and Raltitrexed. **D)** Differential expression analysis for Methotrexate treatment (5µM, 6hrs) for FlO-1, SK-GT-4 and OE33 cell lines. Red indicates genes reaching both P-value and fold change threshold, blue indicates genes that reached p-value threshold and green indicates genes that reached the fold change threshold. P-value = 0.05, log_2_ fold change = 0.5.

Multiparametric phenotypic dose response profiles of the antimetabolites overlaid on the reference library of annotated compounds (**Supplementary Table S2**) show strong dose dependent phenotypic changes, moving from phenotypically inactive (clustering with DMSO controls) to clustering with the DNA damaging agents at active concentrations (**Figure 4B**) in all but the JH-EsoAD1 and MFD-1 lines. All three compounds also showed little or no effect in the control lines EPC2-hTERT and CP-A, clustering closely with the DMSO controls at all concentrations tested.

Class probabilities from the pre-trained machine learning model for each of the compounds predicts them to belong to the DNA damage class for all but the MFD-1 and JH-EsoAD1 lines (**Figure 4C)**, consistent with the clustering above. Probabilities also increase in a dose dependent manner indicating cellular phenotypic activity follows a linear on-target dose response relationship. These results further confirm the ability of the Cell Painting assay to accurately predict MOA of validated hit compounds.

NanoString differential expression analysis revealed methotrexate treatment caused a significant reduction in the expression of Histone H3 subunits (HIST1H3B, HIST1H3G, HISTH3H) (**Figure 4D**) in the sensitive cell lines only, with no effect in either of the tissue matched controls (**Supplementary Table S4)**. Several other genes change with methotrexate treatment but none are significant. Further mechanistic studies shall be required to further elucidate how and if such expression changes confer selectivity to methotrexate.

### Towards novel therapies and targets for OAC

From a subset of 13,000 small molecule compounds with unknown targets we further identify a small number of compound hits from our diverse chemical library which show potent and selective activity for the OAC cell lines. Compound 1 is selective for the OAC-P4C and MFD-1 cells (**Figure 5A**) and machine learning probabilities for all classes are low (**Figure 5B**). Compound 2 induces a strong phenotypic dose response in the OAC-P4C and OE33 cell lines only and does not cluster with the reference library of known MoA (**Figure 5A**). Machine learning predicts it to be DNA damaging (91 % probability) in the OAC-P4C cells, however, its clustering is distinct and the machine learning probability that it is DNA damaging in the OE33 cell line is only 52 % (**Figure 5B**). Therefore it may in fact represent a novel MoA or be acting to cause DNA damage in a novel way. This indicates that these compounds may be selectively targeting novel oesophageal cancer biology. Subsequent transcriptomic and proteomic pathway analysis and target deconvolution studies may reveal the mechanistic pathways involved.

**Figure 5.**
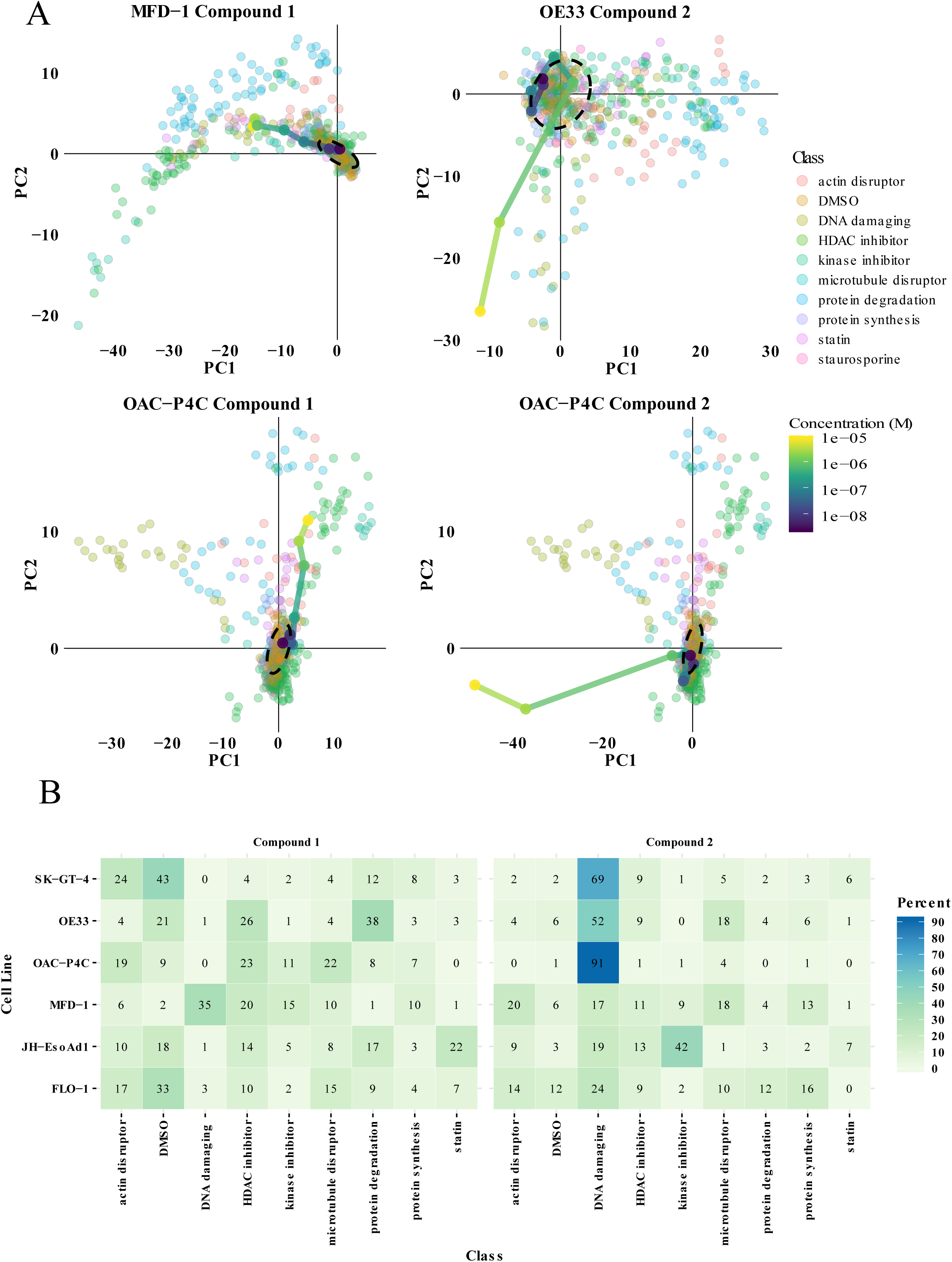
Phenotypic analysis of novel compounds. **A)** Principal component analysis of compound 1 and compound 2 dose responses overlaid on reference library for the two most sensitive cell lines for each compound. **B)** Probabilities expressed as percentages for compound 1 and compound 2 (10µM) belonging to each class in the reference library for each cell line.

## Discussion

Conventional target-directed drug discovery strategies remain to make any impact on the discovery and translation of effective new treatments for oesophageal cancer patients. Key challenges in oesophageal cancer include a highly heterogeneous genetic landscape with few mutations in oncogenic drivers, thereby confounding the identification of clear drug-target hypothesis and modern personalized medicine strategies. In this study we sought to adapt and evaluate the utility of an advanced high-content phenotypic screening method as an empirical strategy for identifying novel drug targets, MoA and pharmacological classes which target OAC.

Here we have shown that combining high-content screening and image informatics with machine-learning can prove effective in the identification and mechanistic characterisation of hit compounds with selective activity upon OAC cell phenotypes. The majority of multiparametric high-content screening assays and associated machine learning methods used to predict drug MoA are typically performed on a single cell line. In this study we have further shown that this format can be applied to heterogeneous panels of cancer cell lines and normal tissue match control cells for the identification and prioritization of hit compounds and MoA which demonstrate selectivity activity for OAC cells.

Machine-learning can be implemented as a tool for multiparametric phenotypic assay quality control (e.g. confirming if the assay is suitable as a discovery platform to classify specific cell phenotypes and elucidate MoA) as well as a tool for MoA deconvolution of hit compounds. Our results demonstrate that this can be standardised across heterogeneous panels of cells with reasonable accuracy.

Following one class of compounds identified in our primary phenotypic screen of 19,555 small molecules tested across all eight oesophageal cell lines, we validated antimetabolites as selectively lethal to the OAC lines *in vitro* following dose response studies. Utilising the multiparametric phenotypic information to generate phenotypic dose responses, combined with a reference library of compounds, machine-learning and clustering techniques we demonstrated the ability to study/predict the MoA of hits from the screen. Here we validated this technique using the antimetabolite hit compounds (Methotrexate, Pemetrexed and Raltitrexed), showing DNA damage as a likely MoA for the selectivity of these compounds which is consistent with the literature ^28,29^. These results together with our identification of hit compounds from our diverse chemical set which are not classified by our reference set of known MoAs demonstrates the impact of phenotypic screening in combination with machine-learning for MoA studies. This strategy will be used to assess and prioritise novel small molecule hits from the diverse chemical library screen for further mechanistic studies. From our primary phenotypic screen we have identified in total 75 compounds which match our hit selection criteria for selective activity across the OAC panel. These 75 hits are an accumulation of the 62 compounds defined by cell morphometric phenotypic analysis and 27 compounds defined by cell proliferation and survival (nuclei count) analysis with 14 compounds overlapping. The 75 hits shall be further progressed through dose-response studies and secondary assays to confirm and prioritize classes of selective compounds for subsequent drug repurposing and or drug discovery studies.

In addition, using bioinformatic approaches we hope that integration of phenotypic data with genetic data across our panel of diverse cell lines may provide insight into the selective activity of phenotypic hits and generate the basis for future genetic biomarker-based clinical trials in OAC.

Overall, our high-content OAC assay has proven effective in the identification and mechanistic characterisation of hit compounds, demonstrating its utility as a novel empirical strategy for the discovery of new therapeutic targets, chemical starting points and repurposing of existing drug classes to re-ignite drug discovery and development in OAC.

## Acknowledgments

This study was supported by an MRC-Institute of Genetics and Molecular Medicine PhD studentship award to REH, the Anne Forrest Fund for Oesophageal Cancer Research and a CRUK Small Molecule Drug Discovery project award to NOC. We also thank Fabrice Turlais and Mathew Calder from CRUK-Therapeutics Discovery Labs for provision of compound libraries and Anil Rustig, University of Pennsylvania for provision of EPC2-hTERT cells.

