## Supplementary Material for "High Content Phenotypic Profiling in Oesophageal Adenocarcinoma Identifies Selectively Active Pharmacological Classes of Drugs for Repurposing and Chemical Starting Points for Novel Drug Discovery"

**Supplementary Table S1. Compound Libraries and screening concentrations.**

| **Library** | **Concentration (μM)** |
|---|---|
| Prestwick Chemical Library | 1 |
| BioAscent 3K Library | 10 |
| LOPAC | 3 |
| Bespoke Library | 1-3 |
| CRUK therapeutics discovery laboratories Library | 10-12 |

**Supplementary Table S2. Reference Library of Compounds**. 38 compounds and their classes.

| **Compound** | **Mechanism of Action** |
|---|---|
| Cytochalastin B | Actin disrupting |
| Cytochalastin D | Actin disrupting |
| Latrunculin | Actin disrupting |
| Camptothesin | DNA damaging |
| SN38 | DNA damaging |
| Dasatinib | Kinase inhibitor |

| | |
|---|---|
| Saracatinib | Kinase inhibitor |
| Epothilone B | Microtubule disrupting |
| Paclitaxel | Microtubule disrupting |
| Colchicine | Microtubule disrupting |
| Nocodazole | Microtubule disrupting |
| Monastrol | Microtubule disrupting |
| ARQ621 | Microtubule disrupting |
| Barasertin | Microtubule disrupting |
| ZM447439 | Microtubule disrupting |
| MG132 | Protein degradation |
| Lacacystin | Protein degradation |
| ALLN | Protein degradation |
| ALLM | Protein degradation |
| Cyclohexamide | Protein synthesis |
| Emetine | Protein synthesis |
| Lovastatin | Statin |
| Simvastatin | Statin |
| SAHA | HDAC inhibitor |
| Panobinistat | HDAC inhibitor |
| Trichostatin A | HDAC inhibitor |
| Romidepsin | HDAC inhibitor |
| Entinostat | HDAC inhibitor |
| Quisinostat | HDAC inhibitor |
| Ricolinostat | HDAC inhibitor |
| Tubastatin A | HDAC inhibitor |
| Droxinostat | HDAC inhibitor |
| PCI34051 | HDAC inhibitor |
| TMP195 | HDAC inhibitor |
| LMK235 | HDAC inhibitor |
| CUDC907 | HDAC inhibitor |
| Belinostat | HDAC inhibitor |
| BR-98 | HDAC inhibitor |

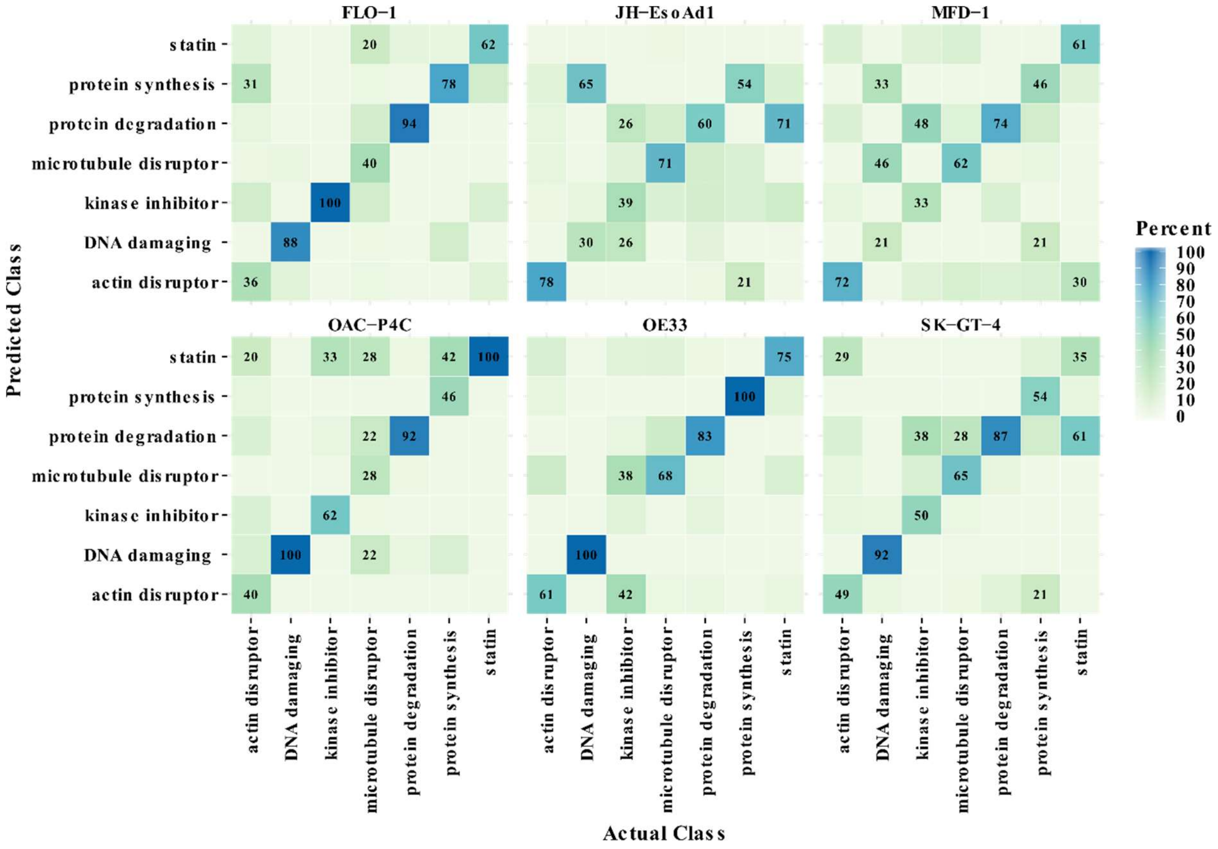


**Supplementary Fig. S1. Leave-one-out random forest confusion matrices for reference library of compounds with known mechanism-of-actions.** Prediction accuracies for each withheld cell line from a random forest classifier trained on the other five cell lines at a time.

A

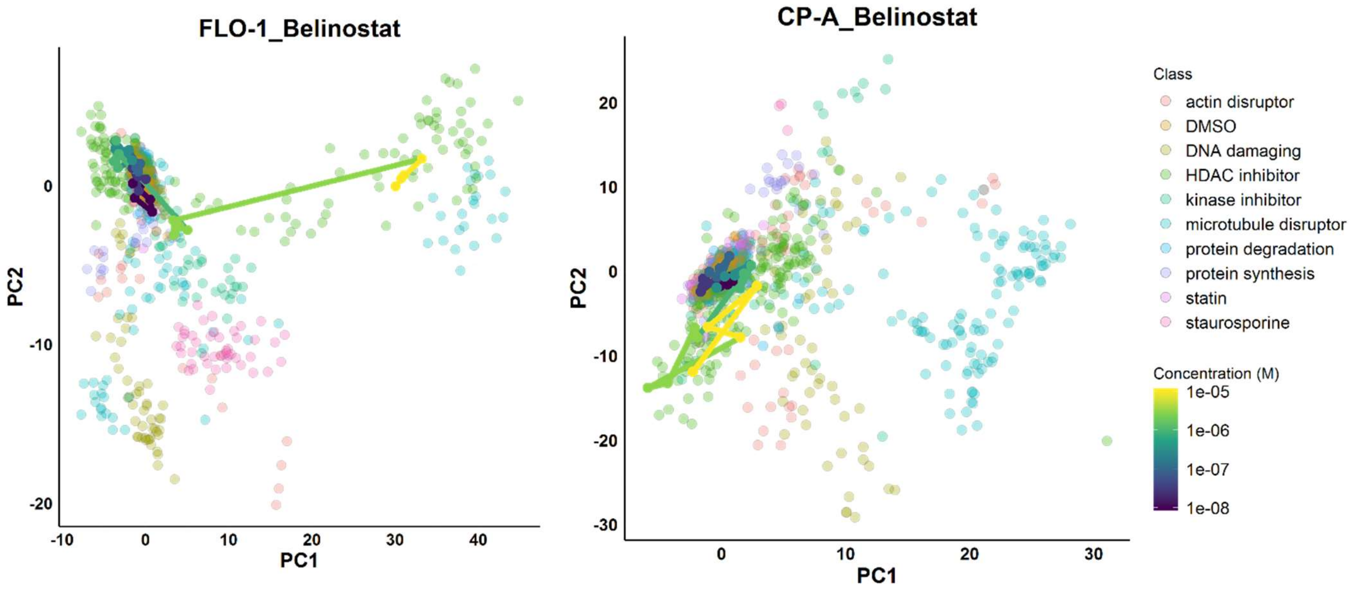


B

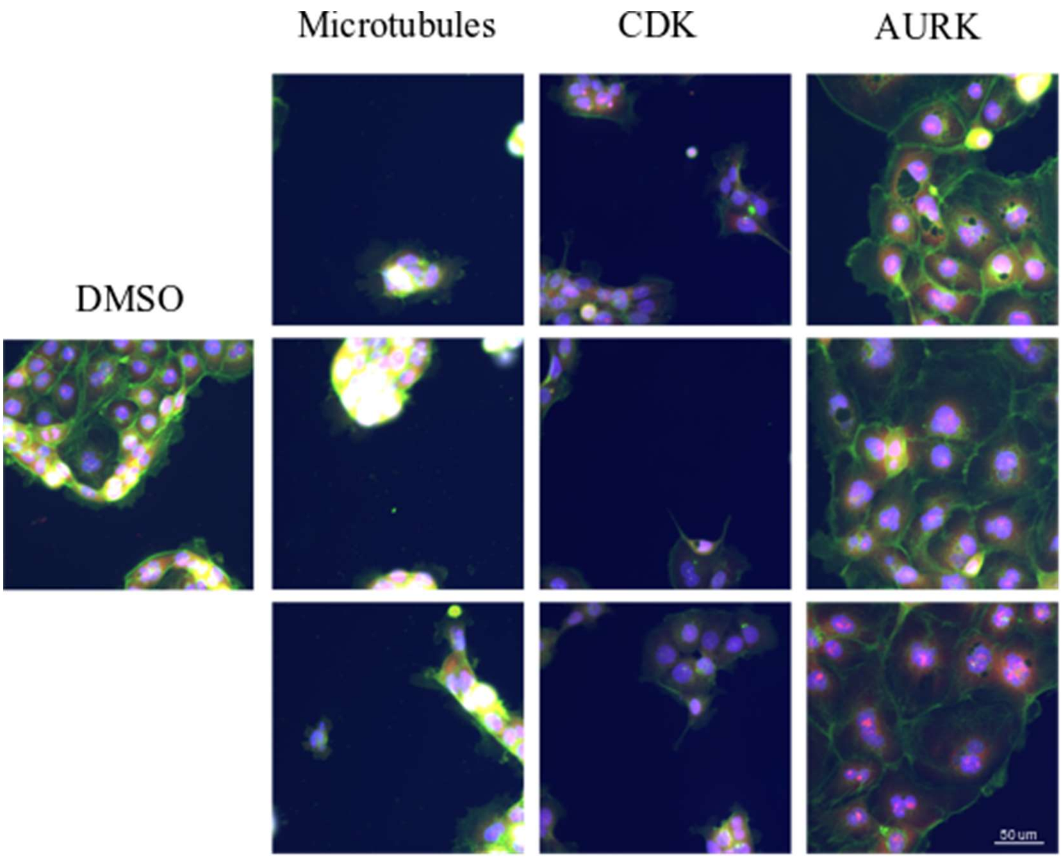


**Supplementary Fig. S2. Phenotypic analysis Data**. A) Phenotypic dose response for HDAC inhibitor Belinostat. The first two principal components for the feature data from the Belinostat dose response overlaid on reference library for FLO-1 and CP-A cell lines. B) Colour combined images for OAC-P4C cells treated with three compounds from each of three classes; Aurora kinase (AURK) inhibitors, cyclin dependent kinase (CDK) inhibitors, Microtubule disruptors (DAPI blue, Phalloidin-TxRED green, Syto$^{TM}$14-CY3 red). Scale bar is 50 μm.

**Supplementary Table S3. Antimetabolite IC50s across the panel of cell lines (nM).**

| Cell Line | Methotrexate | Ralititrexed | Pemetrexed |
|---|---|---|---|
| JH-EsoAD1 | 35 | 2 | 37 |
| FLO-1 | 37 | 4 | 71 |
| MFD-1 | 39 | 8 | 64 |
| OAC-P4C | 49 | 3 | 65 |
| OE33 | 26 | 1 | 25 |
| SK-GT-4 | 20 | 3 | 46 |

**Supplementary Table S4. NanoString normalised counts for Histone H3 subunits**

| | CPA DMSO | CPA MTX | EPC2 DMSO | EPC2 MTX | FLO1 DMSO | FLO1 MTX | OE33 DMSO | OE33 MTX | SKGT4 DMSO | SKGT4 MTX |
|---|---|---|---|---|---|---|---|---|---|---|
| HIST1H3H | 30,333 | 29,853 | 19,088 | 18,171 | 18,179 | 7,925 | 24,946 | 9,563 | 25,135 | 13,495 |
| HIST1H3G | 17,566 | 17,337 | 10,793 | 10,095 | 17,970 | 9,243 | 14,998 | 5,527 | 17,573 | 10,035 |
| HIST1H3B | 26,664 | 25,856 | 17,285 | 16,441 | 28,459 | 18,330 | 30,594 | 15,909 | 27,481 | 17,485 |
